# High-throughput Screening of Compounds against Autoluminescent Nonreplicating *Mycobacterium tuberculosis* under Diverse Conditions

**DOI:** 10.1101/2024.03.10.584296

**Authors:** Xirong Tian, Wanli Ma, Buhari Yusuf, Chunyu Li, H.M. Adnan Hameed, Xinyue Wang, Nanshan Zhong, Jinxing Hu, Tianyu Zhang

**Affiliations:** State Key Laboratory of Respiratory Disease, Guangzhou Institutes of Biomedicine and Health (GIBH), Chinese Academy of Sciences (CAS), Guangzhou 510530, China; Guangdong-Hong Kong-Macao Joint Laboratory of Respiratory Infectious Diseases, Guangzhou Institutes of Biomedicine and Health (GIBH), Chinese Academy of Sciences (CAS), Guangzhou 510530, China; University of Chinese Academy of Sciences (UCAS), Beijing 100049, China; China-New Zealand Joint Laboratory on Biomedicine and Health, Guangzhou Institutes of Biomedicine and Health, Chinese Academy of Sciences, Guangzhou 510530, China; Guangzhou National Laboratory, Guangzhou 510005, China; Guangzhou Chest Hospital, Guangzhou 510005, China

**Keywords:** Autoluminescence, Low-oxygen, Persistence, *Mycobacterium tuberculosis*, High-throughput screening

## Abstract

The screening of new anti-mycobacterial chemicals is primarily focused on inhibiting the active growing bacteria. However, a major challenge in tuberculosis control is the ability of *Mycobacterium tuberculosis* to enter a nonreplicating state for extended periods, rendering it resistant to many clinical drugs and complicating eradication efforts. Existing low-oxygen-recovery assays designed for screening compounds targeting nonreplicating *M. tuberculosis* have limitations, including the colony-forming unit counting for non-luminous *M. tuberculosis* and the instability of the free plasmid carrying *luxAB* genes in luminescent *M. tuberculosis*, along with exogenous substrate requirements for light producing. Moreover, these assays fail to accurately replicate the growth conditions of nonreplicating *M. tuberculosis in vitro*, thus resulting in less convincing results. To address these challenges, we have developed an autoluminescence-based, cholesterol-enriched culture evaluation model to assess 17 anti-tuberculosis drugs of different classes against nonreplicating *M. tuberculosis*. Our findings indicate that the relative light unit, measured in real-time, serves as a reliable surrogate marker for colony-forming unit, which typically becomes available one month later. This suggests the utility of our model for the rapid determination of drug susceptibility dynamically. The autoluminescent *M. tuberculosis*, harbouring *luxCDABE* gene cluster within its genome, can emit blue-green light stably and autonomously without requiring an external substrate supplement. The minimal inhibitory concentrations of all the drugs tested under anaerobic conditions are significantly different from that detected in aerobic environment. Our model allows for rapid, precise, and efficient assessment of drug activity under anaerobic conditions, thereby enabling a more comprehensive evaluation of anti-mycobacterial efficacy. Overall, our model represents a significant advancement in anti-tuberculosis drug discovery and pharmaceutical development.

## INTRODUCTION

The highly infectious pathogen *Mycobacterium tuberculosis* is the causative agent of tuberculosis (TB), and was responsible for 10.6 million TB individuals globally in 2022 (1). The emergence of drug-resistant *M. tuberculosis* strains, coupled with lethal coinfections with human immunodeficiency virus, has significantly exacerbated the global healthcare crisis, necessitating the exploration of novel drugs or treatment regimens against *M. tuberculosis* (2, 3). Notably, after four decades of persistent efforts, bedaquiline (BDQ), the first new anti-TB medication, received approval from the United States Food and Drug Administration in 2012 (4–6).

The persistence of nonreplicating *M. tuberculosis* within lung lesions, capable of surviving for decades and adapting to hypoxic conditions, poses a formidable challenge to TB eradication (7–9). A comprehensive understanding of the complex host environment in which *M. tuberculosis* resides is crucial for developing effective control strategies (10). Previous studies have revealed *M. tuberculosis*’s ability to exploit host-derived cholesterol as a primary carbon source during infection (11). By incorporating cholesterol into growth media, it is more accurately replicate the growth environment of nonreplicating *M. tuberculosis* compared to traditional glycerol-enriched media.

Despite the availability of well-established *in vitro* low-oxygen-recovery assays for assessing evaluate drug efficacy against nonreplicating *M. tuberculosis*, these methods still have certain limitations (12–14). Some methods rely on enumerating colony forming units (CFUs) or visible bacterial growth, requiring several months for the completion of testing. Others have employed a luminescent *M. tuberculosis* strain with a free plasmid carrying *luxAB* genes to screen for chemical agents against nonreplicating mycobacteria (6). However, this approach is not favored due to its dependence on additional substrates and the instability of the plasmid (6). Our research introduces an autoluminescence-based, low-oxygen, and cholesterol-enriched evaluation model that utilizes autoluminescent *M. tuberculosis* H37Ra (AlRa), emitting stable and efficient blue-green light dynamically over an extended period without requiring substrate supplementation. Bacterial growth is quantified dynamically by measuring relative light units (RLUs) at various time points during the recovery process. In conclusion, our model is designed to be straightforward, time-efficient, labor- and resource-effective, while also facilitating high-throughput screening of drugs and drug combinations against nonreplicating *M. tuberculosis*.

## RESULTS

### MICs determination using glycerol- and cholesterol-enriched media under aerobic conditions

Seventeen anti-TB drugs from distinct classes were selected for this study. Cholesterol serves as a primary carbon source during *M. tuberculosis* infection, mimicking the nutritional conditions encountered during the course of *M. tuberculosis* infection. Moreover, under aerobic conditions, the minimal inhibitory concentrations (MICs) of the drugs against *M. tuberculosis* remained largely consistent, irrespective of whether glycerol or cholesterol was used as the carbon source. An exception to this was observed with para-aminosalicylic acid (PAS), as detailed in Table 1 and Figure 1. In the glycerol-enriched medium, the MIC of PAS ranged from 0.25 to 1 μg/mL, however, it increased dramatically to over 64 μg/mL in the cholesterol-enriched medium (Table 1). The growth curves of actively growing AlRa treated with nine drugs from different categories are shown in Figure 1, and those treated with the remaining eight drugs are shown in Figure S1.

**Table 1.** The MICs of clinical drugs against AlRa under aerobic and anaerobic conditions using various media.

| Class/Targets | Drugs * | MIC (µg/mL)/aerobic conditions |  | MIC (µg/mL)/anaerobic conditions |  |
| --- | --- | --- | --- | --- | --- |
|  |  | Glycerol | Cholesterol | Glycerol | Cholesterol |
| Cell Wall | EMB | 2 | 4 | >64 | >64 |
|  | INH | 0.125 | 0.125 | >64 | >64 |
|  | VAN | 16-32 | 16-32 | >64 | >64 |
|  | ETH | 64 | 64 | >64 | >64 |
| RNA polymerase | RIF | 0.0625 | 0.0625-0.125 | 0.125-0.5 | 0.25-2 |
|  | RFB | 0.008 | 0.008 | 0.03125-0.125 | 0.03125-0.0625 |
| Ribosome | AMK | 0.25-1 | 0.25-0.5 | 1-8 | 2-16 |
|  | STR | 0.25-0.5 | 0.25-0.5 | 1-8 | 2-16 |
| Unknown | POA | 400 | 400 | >800 | >800 |
| DNA gyrases | MOX | 0.0625-0.25 | 0.125 | 0.125-2 | 8-32 |
|  | LEV | 0.125-0.5 | 0.25-1 | >64 | >64 |
| Respiration/<br>Production of NO | PA-824 | 0.0625 | 0.0625-0.125 | 1-4 | 1-4 |
| Folic acid synthesis | SMX | > 64 | > 64 | >64 | >64 |
|  | PAS | 0.25-1 | > 64 | >64 | >64 |
| Respiration/<br>Electron transport<br>chain | TB47 | 0.004-0.008 | 0.004-0.008 | 0.125-0.25 | 0.0625-0.25 |
| Respiration/<br>ATP synthesis | BDQ | 0.5-1 | 1-2 | 4-8 | 4-16 |
| Unknown | CLO | 0.5-1 | 1-2 | 2-8 | 8-16 |

\* EMB, ethambutol; INH, isoniazid; VAN, vancomycin; ETH, ethionamide; RIF, rifampin; RFB, rifabutin; AMK, amikacin; STR, streptomycin; MOX, moxifloxacin; LEV, levofloxacin; PA-824, pretomanid; SMX, sulfamethoxazole; PAS, para- aminosalicilic; BDQ, bedaquiline; CLO, clofazimine.

**Figrue 1.**
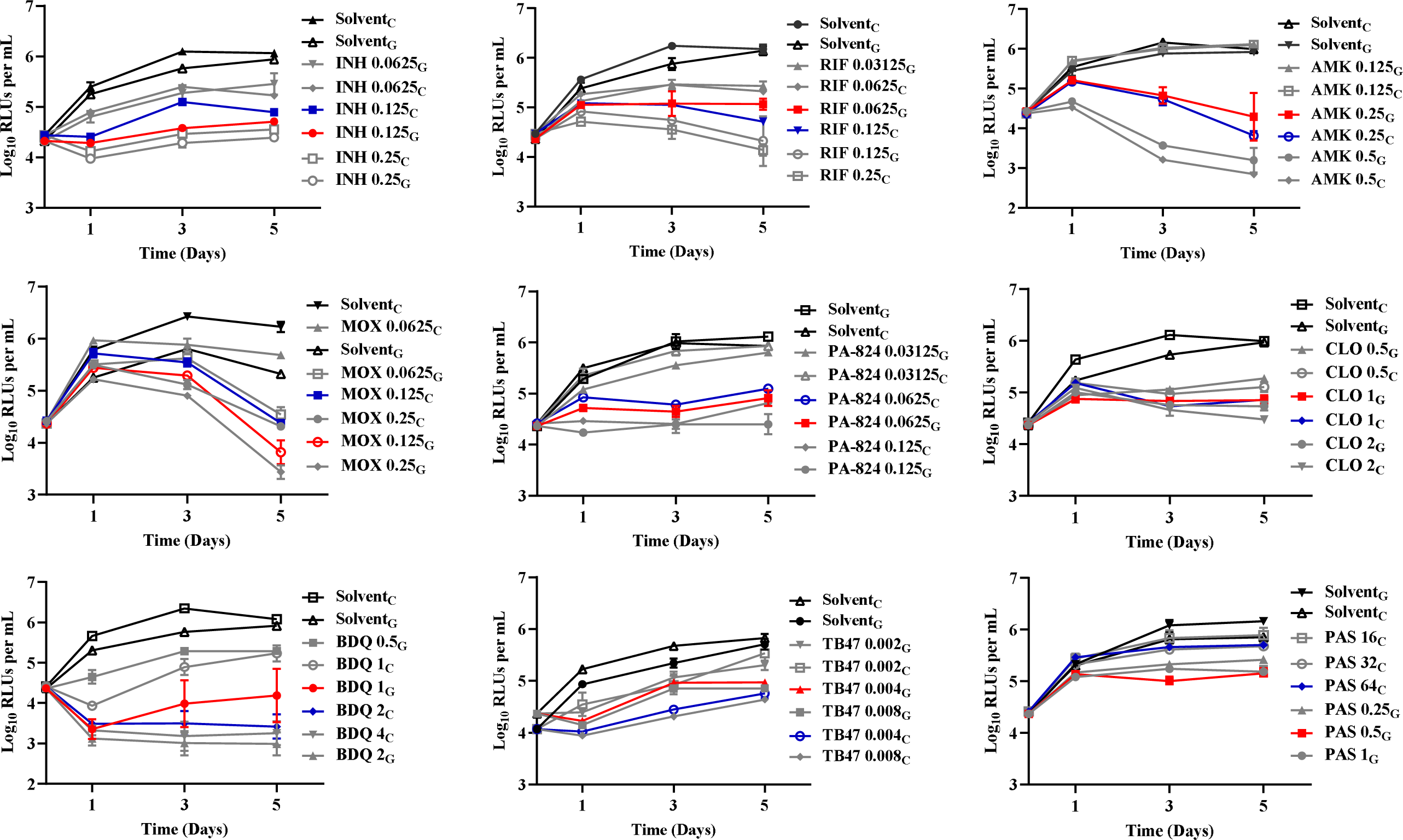
The time-killing curves of actively growing M. *tuberculosis* treated with nine distinct pharmacological categories of drugs aerobic condition using various media. Solvent, DMSO or distilled water; G, 7H9 supplied with glycerol; C, 7H9 enriched with cholesterol; INH, isoniazid; RIF, rifampin; AMK amikacin; MOX, moxifloxacin; PA-824, pretomanid; PAS, para-aminosalicylic; BDQ, bedaquiline; CLO, clofazimine.

### MICs determination using glycerol- and cholesterol-enriched media under anaerobic conditions

We systematically evaluated 17 anti-TB drugs targeting various crucial aspects of *M. tuberculosis* metabolism, including the cell wall, RNA polymerase, ribosome and respiration. Our findings revealed that the MICs of these drugs against *M. tuberculosis* were generally higher under anaerobic conditions compared to aerobic conditions with the exception of rifampin (RIF) and rifabutin (RFB), as depicted in Table 1, Figures 2 and Figure S2. Additionally, the MICs of the drugs remained relatively consistent under anaerobic conditions, regardless of whether the medium was supplemented with glycerol or cholesterol, with the exception of moxifloxacin (MOX). The MICs of MOX against nonreplicating *M. tuberculosis* were found to be in the range of 0.125-2 μg/mL in the presence of glycerol, whereas in the presence of cholesterol, the MICs spanned from 8-32 μg/mL (Table 1). Both MOX and levofloxacin (LEV), the two fluoroquinolones, exhibited increased MICs under anaerobic conditions, regardless of the presence of cholesterol or glycerol in the media. Remarkably, MOX displayed a greater efficacy compared to LEV under both anaerobic and aerobic conditions.

**Figure 2.**
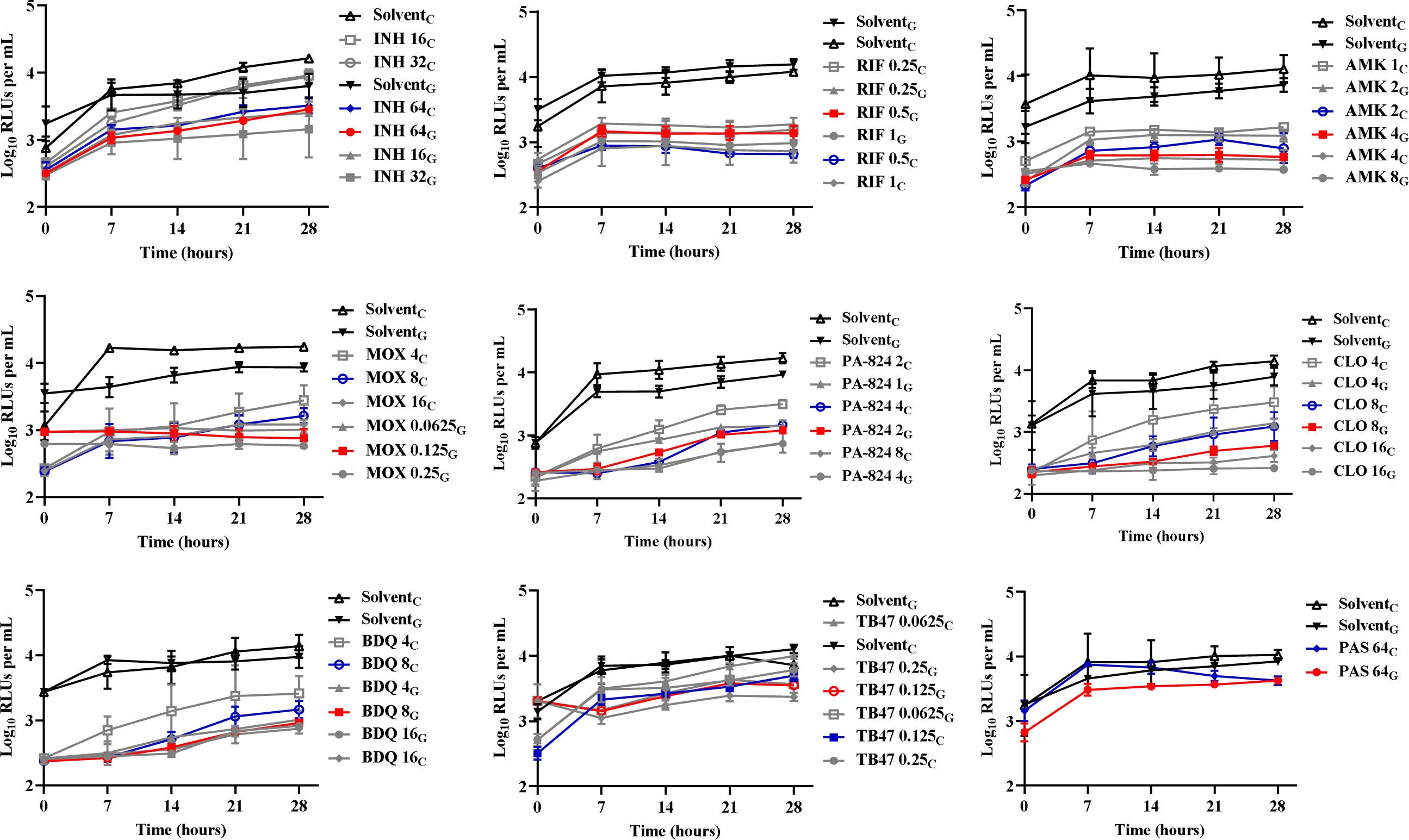
The growth curves of nonreplicating *M. tuberculosis* after treatment with nine distinct pharmacological categories of drugs, various media under anaerobic condition. Solvent, DMSO or distilled water; G, 7H9 supplied with glycerol; C, 7H9 enriched with cholesterol; INH, isoniazid; RIF, rifampin; AMK, amikacin; MOX, moxifloxacin; PA-824, pretomanid; PAS, para-aminosalicylic; BDQ, bedaquiline; CLO, clofazimine.

Under anaerobic conditions, drugs targeting the cell wall, such as vancomycin (VAN), isoniazid (INH), ethambutol (EMB), and ethionamide (ETH), did not display significant antibacterial activity against nonreplicating *M. tuberculosis* as the MICs were over 64 μg/mL (Table 1). In contrast, these drugs exhibited notable efficacy against actively growthing *M. tuberculosis* under aerobic conditions.

Compared to other anti-TB drugs, drugs targeting RNA polymerase showed lower MICs with notably effective anti-*M. tuberculosis* activity in both conditions, particularly under anaerobic conditions. The MICs of RIF ranged from 0.0625 to 0.125 μg/mL and 0.125 to 2 μg/mL under aerobic and anaerobic conditions respectively (Table 1). RFB displayed enhanced efficacy compared to RIF against nonreplicating *M. tuberculosis*, with the MICs as low as 0.03125 to 0.125 μg/mL and 0.03125 to 0.0625 μg/mL in glycerol- and cholesterol-enriched media, respectively (Table 1). These findings emphasized the promising efficacy of RIF and RFB as candidates for eradicating nonreplicating *M. tuberculosis*, thus establishing a robust foundation for their clinical application.

The ribosome-targeting drugs, amikacin (AMK) and streptomycin (STR), demonstrated anti-TB activity against both actively growing and nonreplicating *M. tuberculosis*. However, the MICs against nonreplicating *M. tuberculosis* were significantly higher in comparison to those observed against actively growing *M. tuberculosis*. Pyrazinoic acid (POA), the active form of pyrazinamide, showed no anti-*M. tuberculosis* activity under anaerobic conditions, with the MIC 800 μg/mL. Under anaerobic conditions, drugs targeting folic acid synthesis exhibited no evident anti-*M. tuberculosis* activity. The MIC of PAS ranged from 0.25-1 μg/mL in glycerol-enriched 7H9 medium under aerobic conditions, which significantly increased to over 64 μg/mL in cholesterol-enriched 7H9 medium under both aerobic and anaerobic conditions (Table 1).

Pretomanid (PA-824) demonstrated pronounced anti-*M. tuberculosis* activity, particularly under anaerobic conditions and with cholesterol as the primary carbon source, with a MIC range of 1-4 μg/mL (Table 1). TB47 exhibited remarkable anti-*M. tuberculosis* activity at an exceptionally low concentration of 0.004 μg/mL under aerobic conditions (Table 1). However, its efficacy against *M. tuberculosis* became evident only when the concentration was increased to 0.0625-0.25 μg/mL under anaerobic conditions (Table 1). The MICs of BDQ were 0.5-1 μg/mL and 1-2 μg/mL against actively growing *M. tuberculosis* when cultivated with glycerol or cholesterol, respectively (Table 1). Nevertheless, under anaerobic conditions, these MICs increased to 4-8 μg/mL and 4-16 μg/mL with glycerol or cholesterol, respectively (Table 1). Clofazimine (CLO) exhibited MICs against nonreplicating *M. tuberculosis* ranging from 2-16 μg/mL, significantly higher than those observed against actively growing *M. tuberculosis*, which had an MIC range of 0.5-2 μg/mL (Table 1).

### The combined effect of TB47 and CLO against nonreplicating AlRa

Our results indicated that the combination of TB47 and CLO exhibited partial synergy in terms of sterilizing activity against nonreplicating *M. tuberculosis* under anaerobic conditions, both with glycerol and cholesterol-enriched media with a Fractional Inhibitory Concentration Index (FICI) of 0.75 (Table 2). Notably, the MIC of CLO decreased from 8 to 4 μg/mL in the presence of TB47 at one-fourth of its MIC. These results highlighted the potential of this drug combination for clinical evaluation against nonreplicating *M. tuberculosis*, which has been proved to have strong sterilizing activities against *M. tuberculosis* in a well-established mouse model in two studies carried out by us, and the manuscript of another study is in revision in Antimicrobial Agents and Chemotherapy (15–17). Moreover, the assessment model established in this study for evaluating the efficacy of drug combinations against nonreplicating *M. tuberculosis* showed significant promise for informing future research and testing endeavors in this field.

**Table 2.** MICs of CLO and TB47 against AlRa along with the corresponding interaction profiles of CLO in conjunction with TB47 assessed by checkerboard method under anaerobic condition.

| Drugs<sup>*</sup> | 7H9 + Glycerol |  | 7H9 + Cholesterol |  |
| --- | --- | --- | --- | --- |
|  | MIC<sub>a</sub><sup>#</sup> | MIC<sub>c</sub><sup>#</sup> | MIC<sub>a</sub><sup>#</sup> | MIC<sub>c</sub><sup>#</sup> |
| TB47 | 0.125 | 0.03125 | 0.125 | 0.03125 |
| CLO | 8 | 4 | 8 | 4 |
| FICI<sup>□</sup> / Effect | 0.75 / partially synergistic |  | 0.75 / partially synergistic |  |

<sup>\*</sup>CLO, clofazimine.
<sup>□</sup>FICI, Fractional inhibitory concentration index; The FICI was calculated as the ratio of the MIC of TB47 in combination to the MIC of TB47 alone, plus the MIC of CLO in combination to the MIC of CLO alone. The FICI was calculated to classify the interactions as one of five kinds: synergistic (FICI ≤ 0.5), partially synergistic (0.5 < FICI < 1.0), additive (FICI = 1.0), indifferent (1.0 < FICI ≤ 4.0) or antagonistic (FICI > 4.0) (31).
<sup>#</sup>MIC<sub>a</sub>, MIC alone; MIC<sub>c</sub>, MIC in combination.

## DISCUSSION

During TB infection, *M. tuberculosis* transitions into a nonreplicating state, becoming resistant to many clinical antibiotics and significantly prolonging therapy (18). *M. tuberculosis* can acquire and metabolize essential host-derived lipids, particularly cholesterol, which facilitates disease initiation and progression (19). This necessitates the establishment of a representative model that faithfully replicates the growth conditions of nonreplicating *M. tuberculosis* within the host environment. This model is characterized by oxygen deprivation and cholesterol enrichment of growth medium, with the goal of screening for effective drugs against nonreplicating *M. tuberculosis*. To fulfill this objective, we employed AlRa, a strain preserved in our laboratory, to establish an autoluminescence-based drug efficacy evaluation model for high-throughput screening of compounds against nonreplicating *M. tuberculosis*. The RLUs emitted by AlRa, producing blue-green light, exhibit a strong correlation with CFUs, thus serving as a surrogate marker rather than CFUs for quantifying *M. tuberculosis* numbers in culture (20).

During the establishment of our model, a notable increase in the growth rate of *M. tuberculosis* was observed in cholesterol-enriched medium compared to glycerol-enriched medium, both continuous cultivation under aerobic conditions and recovery after cultivation under anaerobic conditions. This observation supports previous studies suggesting that cholesterol serves as the primary carbon source during *M. tuberculosis* infection (11). Our findings suggested that RIF, RFB, PA-824, and TB47 demonstrated relatively significant potential for inhibiting nonreplicating *M. tuberculosis* under anaerobic conditions, as previously documented (21–23). Conversely, drugs targeting the mycobacterial cell wall, such as INH, EMB, ETH, and VAN, showed negligible anti-mycobacterial activity under anaerobic conditions, consistent with prior published studies (12, 24). The resistance of nonreplicating *M. tuberculosis* to these drugs was attributed to the slowed process of cellular division under anaerobic condition. Drugs that target folic acid synthesis, such as sulfamethoxazole (SMX) and PAS, manifested minimal bactericidal activities against nonreplicating *M. tuberculosis* (Table 1). PAS also showed limited activity against actively growing *M. tuberculosis* with a MIC exceeding 64 μg/mL, when applying the media supplemented with cholesterol. As a prodrug activated through the folate synthesis pathway, PAS can be converted into folate intermediate analogs, thereby disrupting *M. tuberculosis*’s folate metabolism (25, 26). It has been hypothesized that the addition of cholesterol, which prevents PAS from entering *M. tuberculosis*, or the unique mechanism of action of PAS, contributes to its ineffectiveness against *M. tuberculosis* under cholesterol-enriched media. Ribosome-targeting drugs, AMK and STR, exhibited superior activity against nonreplicating *M. tuberculosis* compared to cell wall-targeting drugs (Table 1). The results are consistent with the observations of some Chinese clinicians, who have noted good treatment efficacy in multi-drug resistant TB patients treated with AMK. This suggests that AMK possesses relatively potent activity in controlling *M. tuberculosis* persisters, despite injection is one of its disadvantages. The MIC of POA against nonreplicating *M. tuberculosis* exceeded 800 μg/mL. We speculated that anaerobic environment might influence the membrane potential of *M. tuberculosis*, which may contribute to its resistance to POA (27). Generally, MICs of some drugs against nonreplicating *M. tuberculosis* were generally higher than those against actively growing *M. tuberculosis*, indicating that some drugs clinically effective against actively growing *M. tuberculosis* may not effectively treat nonreplicating *M. tuberculosis* in lung lesions, which is demonstrated clearly in animal work (16, 28). It is noteworthy that fluoroquinolones (either LEV or MOX) have not shown significant effects on persisting *M. tuberculosis* in previous animal studies, which is consistent with the predictions made in this study (16, 28). The combination of TB47 and CLO demonstrated promising partial synergistic activity with a FICI as low as 0.75 under anaerobic conditions (Table 2), which supports the finding observed in the animal studies and is promising for clinical trial (15, 16).

The model developed in this study for assessing drug activity against nonreplicating *M. tuberculosis* possesses several notable advantages. Firstly, this method stimulates the growth conditions of nonreplicating *M. tuberculosis* within the human host by utilizing media supplemented with cholesterol, thereby ensuring maximal fidelity to the natural environment of the pathogen. Secondly, the model offers simple manipulation as AlRa consistently emits blue-green light without requiring additional substrate aldehyde, thereby guaranteeing stable and consistent results. Thirdly, the results are easily visualized through the consistent detection and comparison of RLUs emitted by AlRa at various time intervals. Further exploration should prioritize high-throughput screening of prospective compounds and treatment regimens against nonreplicating *M. tuberculosis* using the current model. In summary, this model holds significant application value and possesses high potential to accelerate the discovery and development of new drugs and treatment strategies targeting nonreplicating *M. tuberculosis*, which may closely mirror the real state of *M. tuberculosis* persisters *in vivo*.

## MATERIALS AND METHODS

### Mycobacterial strains and culture conditions

The mycobacterial strain utilized in this study is AlRa, an autoluminescent *M. tuberculosis* H37Ra, which has been genetically engineered to incorporate the *luxCDABE* gene cluster into its genome (20). AlRa emits blue-green light continuously catalyzed by LuxAB (bacterial luciferase) and facilitated by LuxCDE (fatty acid reductase complex), which consistently supply the necessary substrate aldehyde in a recycling form, thereby obviating the need for external substrate supplementation (20). Notably, the drug susceptibility and growth rate of AlRa do not significantly differ from those of the wild type *M. tuberculosis* H37Ra (20). AlRa was cultured at 37L in Middlebrook 7H9 broth (Difco, Detroit, MI, USA) supplemented with 10% oleic acid-albumin-dextrose-catalase enrichment (BBL, Sparks, MD, USA), 0.2% glycerol, and 0.05% Tween 80.

### Antimicrobials

All pharmaceutical drugs, including ETH, INH, VAN, RIF, RFB, AMK, STR, MOX, LEV, PA-824, SMX, PAS, POA, BDQ, and CLO, were purchased from Meilun (Dalian, China) with a minimum purity threshold of ≥ 95%. Additionally, TB47 was synthesized and sent to us as a gift by Boji (Guangzhou, China). AMK, INH, and STR were solubilized in sterilized water, while the remaining drugs were dissolved in dimethyl sulfoxide (DMSO) obtained from Xilong (Shanghai, China). All drug solutions were freshly prepared and stored at 4L until use.

### Chemotherapy under aerobic condition

The AlRa were cultured in 7H9 medium supplemented with Tween80 at 37L until the optical density, measured at a wavelength of 600 nm, reached to 0.6-0.8 and the RLUs/mL reached 5 million. The various drug solutions (4 μL each) were combined with AlRa culture (196 μL, 10-fold diluted using 7H9 without Tween80 and supplemented with either glycerol or cholesterol) in a sterilized tube and mixed thoroughly. Negative controls were prepared using DMSO and sterilized water. Each drug concentration and negative control were assayed simultaneously in three 1.5 mL EP tubes. The AlRa-drug mixtures underwent a five-day incubation period, during which the RLUs were consistently assessed at identical intervals each day. The MIC_lux_ (MIC based on the detection of autoluminescence) of drugs against the actively growing AlRa is defined as the lowest concentration at which the RLUs in the experimental group are equal to or less than 10% of the RLUs in the control group after 5 days.

### Culture of nonreplicating AlRa

The AlRa strains were cultured until they reached an optical density of 0.6-0.8, measured at a wavelength of 600 nm, and a state of 5×10^6^ RLUs/mL. Subsequently, the AlRa culture was then transferred to a low-oxygen cabinet (HUAPO, Guangzhou, China) that had already achieved a steady state. This cabinet was programmed to maintain specific conditions, including a temperature of 37L, an oxygen content of 0.8%, and a carbon dioxide concentration of 5%. The methylene blue solution was then added to the AlRa culture to achieve a final concentration of 6 μg/mL (29, 30). The cultures were incubated until the blue color disappeared, typically observed within 7-10 days after the addition of methylene blue.

### Drug susceptibility tested under anaerobic condition

Under hypoxic conditions, various drug solutions (4 μL each) were combined with nonreplicating AlRa culture (196 μL, 10-fold diluted using 7H9 without Tween80 and supplemented with glycerol or cholesterol) in a sterile tube and mixed thoroughly. The AlRa-drug mixture was then incubated for seven days. To remove any potential carryover effects of residual drugs during the subsequent recovery process, activated carbon (in a volume ratio of 1:5, 50 μL into 200 μL) was added (15). The tubes were subsequently transferred to a standard aerobic incubator. The RLUs in the co-incubated 1.5 mL EP tubes were measured and recorded at 7-hour intervals over a period of 28 hours. The experimental procedure was repeated three times.

### Combined activity assay of TB47 and CLO against nonreplicating AlRa

Previous research from our group has demonstrated that the combined administration of TB47 and CLO significantly exhibits anti-replicating *M. tuberculosis* activity both *in vitro* and *in vivo* (16). In this study, we aim to determine whether this combination would also display notable efficacy against nonreplicating *M. tuberculosis* using the checkerboard method. Specifically, 2 μL of TB47, 2 μL of CLO, and 196 μL of diluted AlRa culture in a nonreplicating state were simultaneously added to the same 1.5 mL tube. To eliminate any potential carryover effects of the drugs following seven days of coincubation, activated carbon (at a volume ratio of 1:5, 50 μL into 200 μL) was utilized (26). Subsequently, the tubes containing the co-culture were transferred to aerobic conditions, and RLUs were then detected at 7-hour intervals for the following 28 hours. The FICI was calculated as the ratio of the MIC of TB47 in combination to the MIC of TB47 alone, plus the MIC of CLO in combination to the MIC of CLO alone (31). The effects were categorized based on the FICI into five groups: synergistic (FICI ≤ 0.5), partially synergistic (0.5 < FICI < 1.0), additive (FICI = 1.0), irrelevant (1.0 < FICI ≤ 4.0), and antagonistic (FICI > 4.0) (31).

### Statistical analysis

Before performing statistical analysis, the RLU values were transformed into their logarithmic counts. The time-kill curves were graphically depicted using GraphPad Prism version 8.3.0.

## ACKNOWLEDGMENTS

This work received support from the National Key R&D Program of China (2021YFA1300904), the National Natural Science Foundation of China (21920102003) and by the Guangzhou Science and Technology Planning Project (2023A03J0992). The funders had no role in study design, data collection and analysis, decision to publish, or preparation of the manuscript.

## AUTHOR CONTRIBUTIONS

Conceived the project: Tianyu Zhang, Xirong Tian, Wanli Ma, Nanshan Zhong; Designed the research: Tianyu Zhang, Xirong Tian, Wanli Ma, Xinyue Wang, Jinxing Hu, Nanshan Zhong; Performed the studies: Xirong Tian, Wanli Ma, Chunyu Li, Xinyue Wang; Interpreted the results: Tianyu Zhang, Xirong Tian, Wanli Ma, Jinxing Hu; Drafted the manuscript: Tianyu Zhang, Xirong Tian, Wanli Ma, Buhari Yusuf, H.M. Adnan Hameed; Final approval of the manuscript: all authors.

