## Supplementary material for "High-throughput Screening of Compounds against Autoluminescent Nonreplicating *Mycobacterium tuberculosis* under Diverse Conditions": Figure S1 and Figure S2

**Supplementary materials**

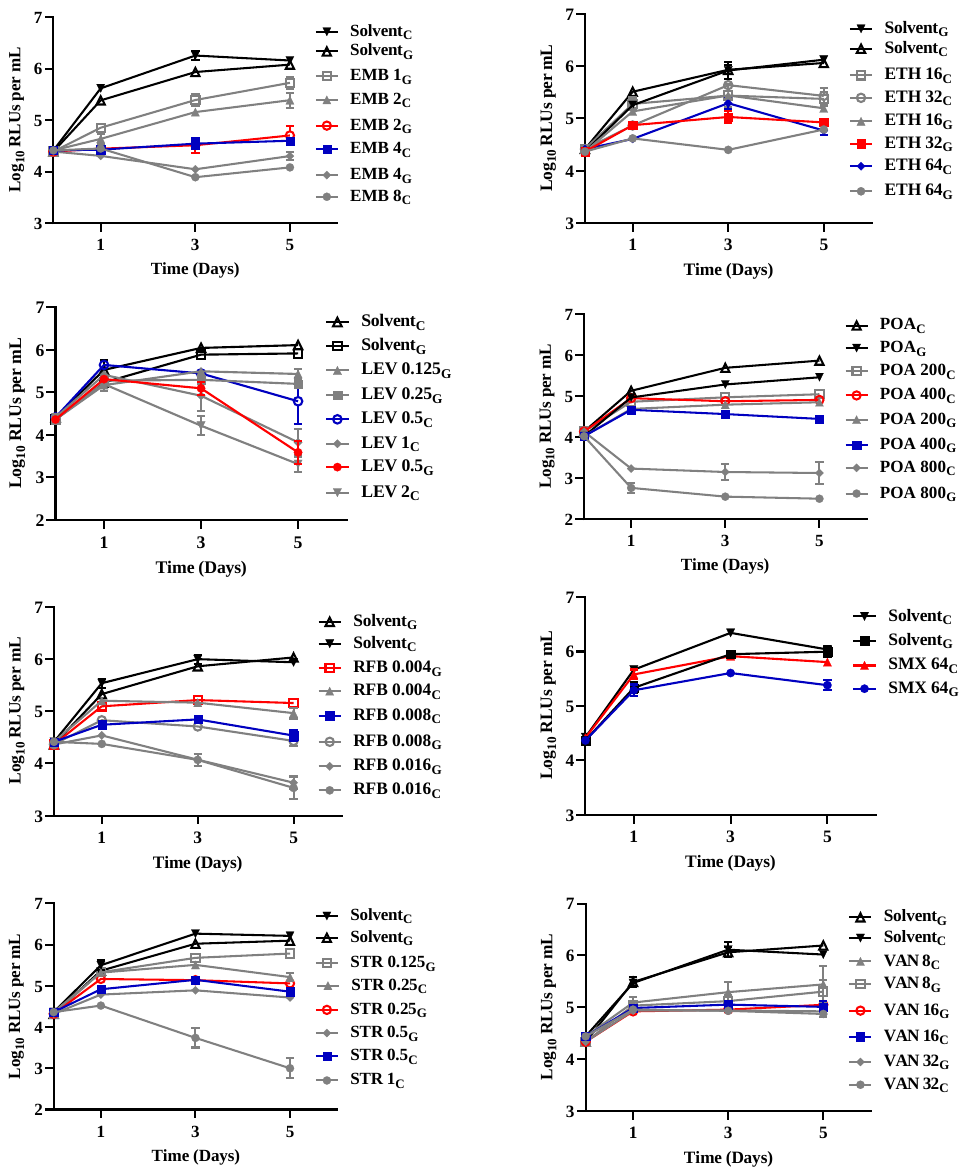

**Figure S1. Time-killing curves of actively growing *M. tuberculosis* treated with eight distinct categories of drugs under aerobic conditions using various media.** Solvent, DMSO or distilled water; G, 7H9 enriched with glycerol; C, 7H9 enriched with cholesterol; EMB, ethambutol; ETH, ethionamide; LEV, levofloxacin; POA, pyrazinoic acid; RFB, rifabutin; SMX, sulfamethoxazole; STR, streptomycin; VAN, vancomycin.

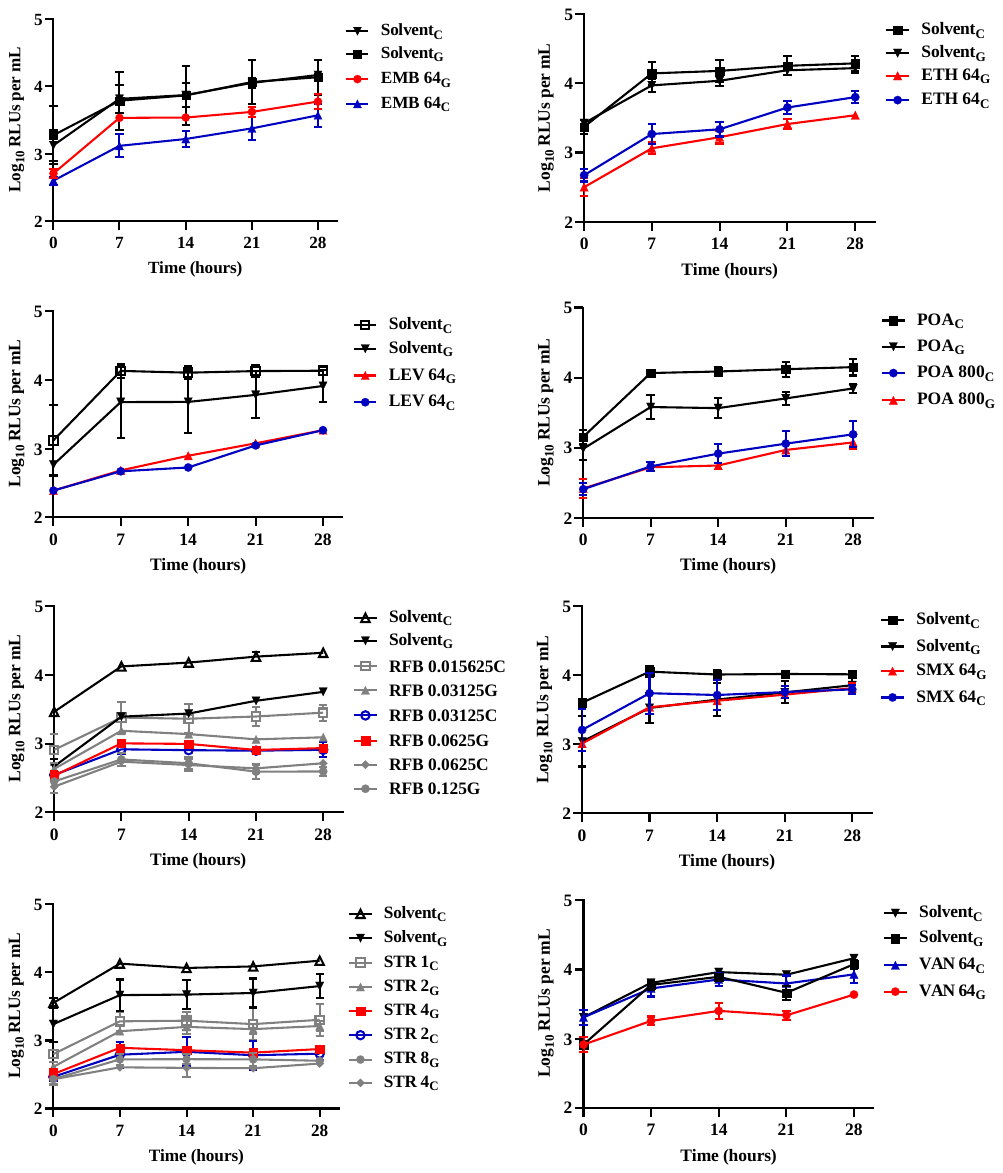

**Figure S2. Growth curves of non-replicating *M. tuberculosis* after treatment with eight different categories of drugs, using various media under anaerobic conditions.** Solvent, DMSO or distilled water; G, 7H9 enriched with glycerol; C, 7H9 enriched with cholesterol; EMB, ethambutol; ETH, ethionamide; LEV, levofloxacin; POA, pyrazinoic acid; RFB, rifabutin; SMX, sulfamethoxazole; STR, streptomycin; VAN, vancomycin.
